# Experience does not change the importance of wind support for migratory route selection by a soaring bird

**DOI:** 10.1101/2022.03.08.483440

**Authors:** Hester Brønnvik, Kamran Safi, Wouter M. G. Vansteelant, Patrik Byholm, Elham Nourani

**Affiliations:** Department of Migration, Max Planck Institute of Animal Behavior, Radolfzell, Germany; Department of Biology, University of Konstanz, Konstanz, Germany; Department of Wetland Ecology, Estación Biológica de Doñana, Spain; Theoretical and Computational Ecology, Institute for Biodiversity and Ecosystem Dynamics, University of Amsterdam, Amsterdam, The Netherlands; Novia University of Applied Sciences, Ekenäs, Finland; Organismal & Evolutionary Biology, University of Helsinki, Finland

**Keywords:** behavioral development, ontogeny, bird migration, step selection function, *Pernis apivorus*

## Abstract

Migration is a complex behavior that is costly in terms of time, energy, and risk of mortality. Thermal soaring birds rely on airflow, specifically wind support and uplift, to offset their energetic costs of flight. Their migratory routes are a record of movement decisions to negotiate the atmospheric environment and achieve efficiency. Because thermal soaring is a complex flight type that young birds need to learn, we expected that, as individuals gain more experience, their movement decisions will also increasingly favor the best thermal uplift conditions. We quantified how route choice during autumn migration of young European honey buzzards (*Pernis apivorus*) was adjusted to wind support and uplift over up to four years of migration and compared this to the choices of adult birds. We found that wind support was important in all migrations. However, we did not find an increase in the use of thermal uplifts, which could be an artifact of the uplift proxies that we used. Age-specific variations in response to airflow might occur at a smaller spatio-temporal scale than we investigated.

## 1 Introduction

Billions of animals migrate, engaging in a challenging behavior during which environmental conditions affect fitness through survival and breeding success [1, 2, 3]. Migrating birds move through the air, which is in motion itself. The most important way for them to offset the energetic costs of movement is to ride airflow. Winds subsidize flight costs when birds move in the same direction as them (wind support), but increase costs of flight when flowing in the opposing direction (headwinds) or perpendicular to the birds’ headings (crosswinds) [4, 5]. Birds can also take advantage of rising air (uplift) [6, 7, 8], whereas sinking air (subsidence) forces them to use powered flight to maintain altitude [9].

Thermal soaring birds are among the most dependent on the dynamics of airflow due to their prohibitively high energetic costs of powered flight [10]. Under optimality theory [11, 12], soaring birds will respond to the costs and benefits of airflow by optimizing their travel within it—by taking advantage of wind support and uplift while avoiding headwinds and crosswinds. This is reflected in their small-scale movement decisions, which determine the energetic costs of larger scale movement behaviors, such as migration [13]. The migratory routes of these birds should be characterized by wind support [14, 4] and uplift [15, 10] to be optimal. Yet not all individuals perform optimally. In many species migratory routes vary in space and time within and between individuals [16, 17], with first-year migrants performing less optimally than more experienced birds [18, 19, 20, 21].

Whether these differences between juveniles and adults develop through a continuous process or a rapid acquisition of behavior remains an important question in behavioral ecology [22, 23, 24]. Here we address this question by investigating how individuals’ improvements in the use of airflow enable them to optimize their migration route choice with experience. We expect that juvenile soaring birds are able to use wind from an early age [25], as flying with wind support is not a cognitively complex task [26]. In contrast, soaring flight is complex, requiring the integration of cognitive processes (perception of the environment to locate thermals) and motor skills (adjusting flight speed and body angle within thermals) and is learned and perfected over time [6]. We therefore expect that younger migrants are not able to take advantage of thermals as efficiently as adults and that their small-scale movement decisions during migration reflect this.

We use a long-term data set of GPS-tracked European honey buzzards to compare the influence of airflow on route choice during successive migrations by juvenile birds. Juvenile and adult honey buzzards differ in their migratory timing and routes. Adults depart the breeding grounds sooner [27] and make long detours around water bodies [28]. Juveniles depart after adults, which leaves them unable to learn from informed conspecifics. They move with prevailing winds, apparently using compass direction and wind to determine their routes [25] and are more likely to cross the sea [28].

We expect the adult behavior to represent an attempt at optimality, and thus that as young birds gain experience their responses approach those of the adults. We expect that (i) wind support is an important determinant of route selection regardless of experience, and (ii) birds increasingly select their routes on the basis of uplift availability as they age. Finally, wind support and uplift are not mutually exclusive, and soaring birds can select routes by responding to one based on the condition of the other [29]. We expect that (iii) inexperienced juveniles are limited to using uplift when wind support is favorable, whereas experienced birds maximize uplift regardless of wind support conditions.

## 2 Methods

### 2.1 Study system

We used existing data from a study of honey buzzards breeding in southern Finland (for details see Vansteelant et al. [25]). Between 2011 and 2014, nine adults of unknown age and 28 fledglings were equipped with Argos or GPS transmitters; they were then tracked for up to seven years. We analyzed the routes taken on autumn migrations so that we could compare the first migration to subsequent journeys.

Five of the fledglings suffered mortality or tag failure before completing their first migrations and were removed from the analysis. For the remaining 23 fledglings, we analyzed the autumn migrations from Finland to sub-Saharan Africa (electronic supplementary material, S1). In addition, five adults of unknown age transmitted four or five autumn migrations. We analyzed the fourth and fifth transmitted routes of these five adults because they are at least the fifth and sixth migrations (after at least one migration in juvenile plumage).

### 2.2 Step-selection functions

#### 2.2.1 Track processing

The transmitters had different sampling rates, ranging from median one to four hours. We re-sampled the tracks of each individual to match the median sampling rate of its transmitter so that time intervals between locations were consistent within each individual across years.

We analyzed route selection using step-selection functions [30, 31], which model movement as a series of discrete steps between consecutive locations, comparing conditions at locations that the birds used to those that were available but forgone.

We generated a stratified data set (electronic supplementary material, S2) for the step selection analysis. For each step along the migratory route (i.e. used step), we generated 100 available steps. We determined the locations of available steps by randomly sampling from gamma distributions fitted to the step lengths and von Mises distributions fitted to the turn angles in the empirical data for each track (using the “amt” package [32] in R [33]; electronic supplementary material, S3). This choice of turn angles accounted for the birds’ direction of migration, as most of the available points were in this direction. Each set of alternative steps also included points in other directions. This distribution of available points provided the model with samples of the range of conditions available to the animals, preventing overestimation of the impact of wind support.

#### 2.2.2 Environmental data

We annotated all used and alternative locations using the Movebank Env-DATA service [34] to obtain data from the European Centre for Medium-Range Weather Forecasts (ECMWF) Global Atmospheric Reanalysis (ERA5). We considered two different proxies for uplift that are available through ECMWF: vertical velocity of pressure (Pa/s), which quantifies vertical air movement, and planetary boundary layer height (m), which is dependent on rising air and therefore is higher where thermal uplift is strong. Both of these variables have been used by previous studies as proxies for uplift strength [35, 36, 37]. For each location we retrieved east/west and north/south wind velocities (m/s), vertical velocity, and planetary boundary layer height. Because pressure is lower with increasing altitude, negative vertical velocity values indicate uplift [38]. All predictors are measured hourly at 0.25 degree (roughly 30 km) resolution and velocities are linearly interpolated at 925 mB pressure level (roughly 762 m asl). We calculated wind support along each used and available step using the east/west and north/south wind velocities [39].

#### 2.2.3 Model fitting

We estimated step selection functions using the integrated nested Laplace approximation (INLA) method (using the “INLA” package [40] in R v. 4.0.2 [33]). We were interested in the importance of wind support, uplift, and the interaction of the two to route selection and whether experience influenced the importance of these variables. We therefore included a three-way interaction term of uplift, wind support and migration year as our predictor. Migration year was included as a continuous variable. All adult birds of unknown age were assigned to migration year 5. We did not find a correlation between our two uplift proxies (vertical velocity and boundary layer height; *r* = -0.11; *p* < 0.05). Thus, We built separate models using the two uplift proxies, Model A with boundary layer height and Model B with vertical velocity as the proxy for uplift. To make the coefficients of our models comparable, we standardized the predictor variables across the whole data set (by calculating z-scores). In each model, we included individual ID as a random effect on the slopes. We set priors of N(0, 10^−4^) for fixed effects and set penalized complexity priors of PC(3, 0.05) to the precisions of the random slopes [41]. Finally, we assessed model fit using conditional predictive ordinates (CPO) and marginal likelihood (MLik). CPO is the probability of detecting a given observation if the model is fit excluding that observation, thus CPO detects outliers. High values of CPO are considered to show good predictive ability. Marginal likelihood is the joint probability of the data averaged over the prior. Smaller values of marginal likelihood are considered to show better fit [42]. We used CPO and MLik to compare the performance of the two models to decide which uplift proxy to use for interpreting the results.

## 3 Results

Due to high juvenile mortality and/or tag failure, of our 23 first-time migrants only five transmitted a second autumn migration, four a third, and just two transmitted a fourth autumn migration. The realized atmospheric conditions along the routes of individuals that transmitted multiple autumn migrations did vary over time (Fig. 1), but on average did not differ among migrations (electronic supplementary material, S4).

**Figure 1:**
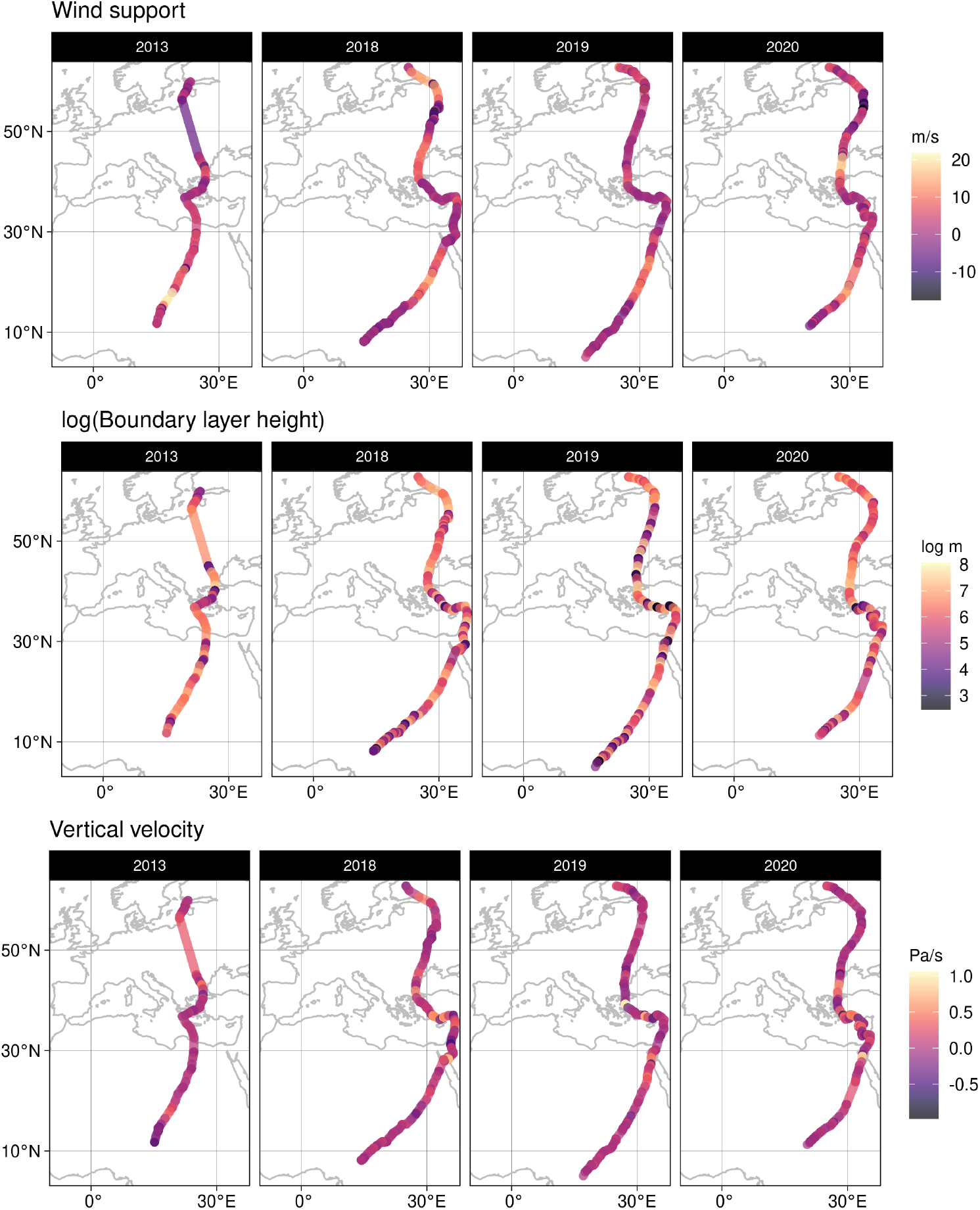
The tracks of a single individual tracked for four autumn migrations (columns left to right). Each track is labeled with the atmospheric conditions that were predictors in our models (rows).

The most important variable predicting route selection was wind support in both models (Fig. 2. In Model A (including boundary layer height), both wind support and the interaction of wind support with uplift were positive and important. In Model B (vertical velocity), only the effect of wind support was important. Migration year (one to five) was not an important predictor of route selection in either model. Models A and B performed equally (CPO = 0.97 for each, Mlik = -38517.73 and -38534.20 respectively). We detected some non-significant individual variation in the wind support and uplift coefficients (electronic supplementary material, S5).

**Figure 2:**
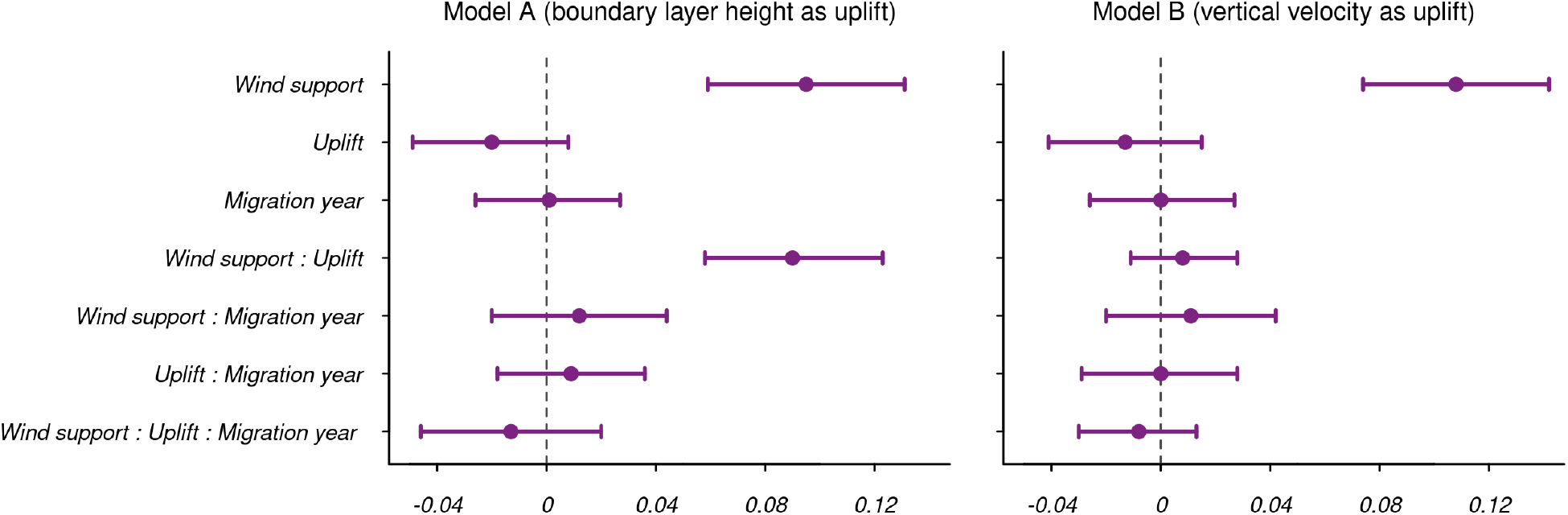
The importance of wind support, uplift, and their interaction to route selection over migrations. Posterior means (centred and scaled) and 95% credible intervals for the fixed effects in the INLA models are shown.

## 4 Discussion

We found that even young European honey buzzards exploit airflow when selecting their migratory routes and that there are no discernible differences between ages. We expected the importance of wind support for route selection of all ages [25, 14, 4], as flying with supportive wind is a cognitively cheap way of reducing flight costs [26]. Contrary to our expectation, there was no variation in responding to uplift among ages.

The role of uplift in route selection was inconsistent in our results. We used two different proxies for uplift, boundary layer height and the vertical velocity of pressure. Neither proxy was significant in our models. However, when we used boundary layer height, the interaction between uplift and wind support was shown to influence route selection positively. Based on these results we cannot make conclusions about the role of uplift in route selection in any age group. The fact that the two proxies were not correlated highlights that the results of studies that investigate soaring bird ecology using such proxies may not be comparable.

Regardless of the proxy used, that we did not find any importance of uplift or an effect of age on route selection may be a matter of scale. The atmospheric data measured at roughly 30 km every hour might have failed to capture the uplift conditions that the birds select, which are likely on the scale of tens of meters. We examined selection at the scale of entire migratory routes, but birds may make adjustments within and between thermals that we could not see, and capacity for these fine scale adjustments may differ between ages [13, 6]. Thus, weather models may not allow us to see the fine-scale improvements birds make after learning to soar and while they learn to soar efficiently. Proxies retrieved from weather models are not as reliable for measuring the proportion of a route spent soaring as data extracted from animals’ movement are. Capturing this movement requires high temporal-resolution GPS data to show circling flight [43] and/or tri-axial accelerometry data to show flapping bouts [44]. These are not currently available for the honey buzzards.

To construct the developmental trajectories in selecting the optimal migratory route, we used a unique data set that allowed us to compare the behavior of the same individuals over up to four years of migration with individuals tagged as adults. Yet the lack of variation in the behaviors among the different ages could indicate that the learning period is longer than 4-5 years. The birds tagged as adults in our study were individuals of unknown ages—it may be that they were still immature and had not yet attained the optimal, adult-like behavior that we expected. Selecting optimal migratory routes may be cognitively demanding, requiring adequate perception of and responses to a changing environment. Thus, improvement in using airflow to optimize soaring flight and migratory performance may be slow and gradual [22, 6, 24]. In long-lived species such as the European honey buzzard, individuals may spend years acquiring and then refining their flight skills and migratory route selection [45]. Thus, data collected for many individuals over extended time periods are required to understand how complex, cost-saving behaviors develop. Our data may simply not be of sufficient duration and resolution for capturing these fine-scale improvements.

Route selection behavior is more complex than simply reducing local energy expenditure, which was the basis of our expectations. Migratory decisions from departure time and travel speed to which routes to use are affected by many factors. Time is important among these as a currency governing migratory decisions along with energy; optimal migration is a compromise between minimizing time and maximizing energy gain [46, 47]. In addition, factors such as predator avoidance [48], availability of roost sites [49] and food [50], and extreme conditions [51, 1, 52] contribute to migratory decisions in ways not considered here and that may differ between ages.

Migration is a complex behavior that can be improved by experience [22, 20, 24]. We studied the ontogeny of migratory route selection in a long-lived, long-distance, soaring migrant in relation to airflow. We show that European honey buzzards use wind support to select migratory routes and that this does not change with experience. Our finding suggests that wind support is important for migration in all life stages of this species and we suspect that this may be the case in other obligate soaring species as well. This may have consequences for the longevity of the species in the face of shifting wind patterns driven by anthropogenic global warming.

## Acknowledgements

We thank the members of the Animal-environment Interactions research group at the Max Planck Institute of Animal Behavior for their valuable comments and discussions throughout this study. We thank M. Honkiniemi, A. Rossi, A. Rantamäki, J. Valkama, J. Kivelä, I. Nousiainen, K. Palo and M. Lehtonen for assisting with fieldwork in Finland. We also thank three anonymous reviewers for their comments and suggestions.

## Ethics

Capturing of the birds was done under a ringing permit (permit 2604) issued by the Finnish Museum of Natural History. Attachment of tracking devices was permitted under the following licenses issued by Finnish authorities: EPOELY/135/07.01.2013, ESAVI/2195/04.10.07/2014, PIRELY/49/07.01/2013, VARELY/73/07.01/2013, VARELY/215/2015.

## Data Accessibility

Data and R scripts used for step selection and permutation analysis are available as electronic supplementary material.

## Authors’ Contributions

EN conceived the study and supervised the project. PB provided the data. WMGV assisted with data preparation. HB analyzed the data with input from KS and EN. HB wrote the first draft of the manuscript. All authors discussed the final results and commented on the manuscript.

## Competing Interests

We declare we have no competing interests.

## Funding

Financial support for the fieldwork was provided by Kone Foundation, Swedish Cultural Foundation in Finland, R. Erik Serlachius Foundation and Aktia Foundation (all to PB). EN was supported by the PRIME programme of the German Academic Exchange Service (DAAD) with funds from the German Federal Ministry of Education and Research (BMBF).

## Supplemental materials

**Supplemental material 1:**
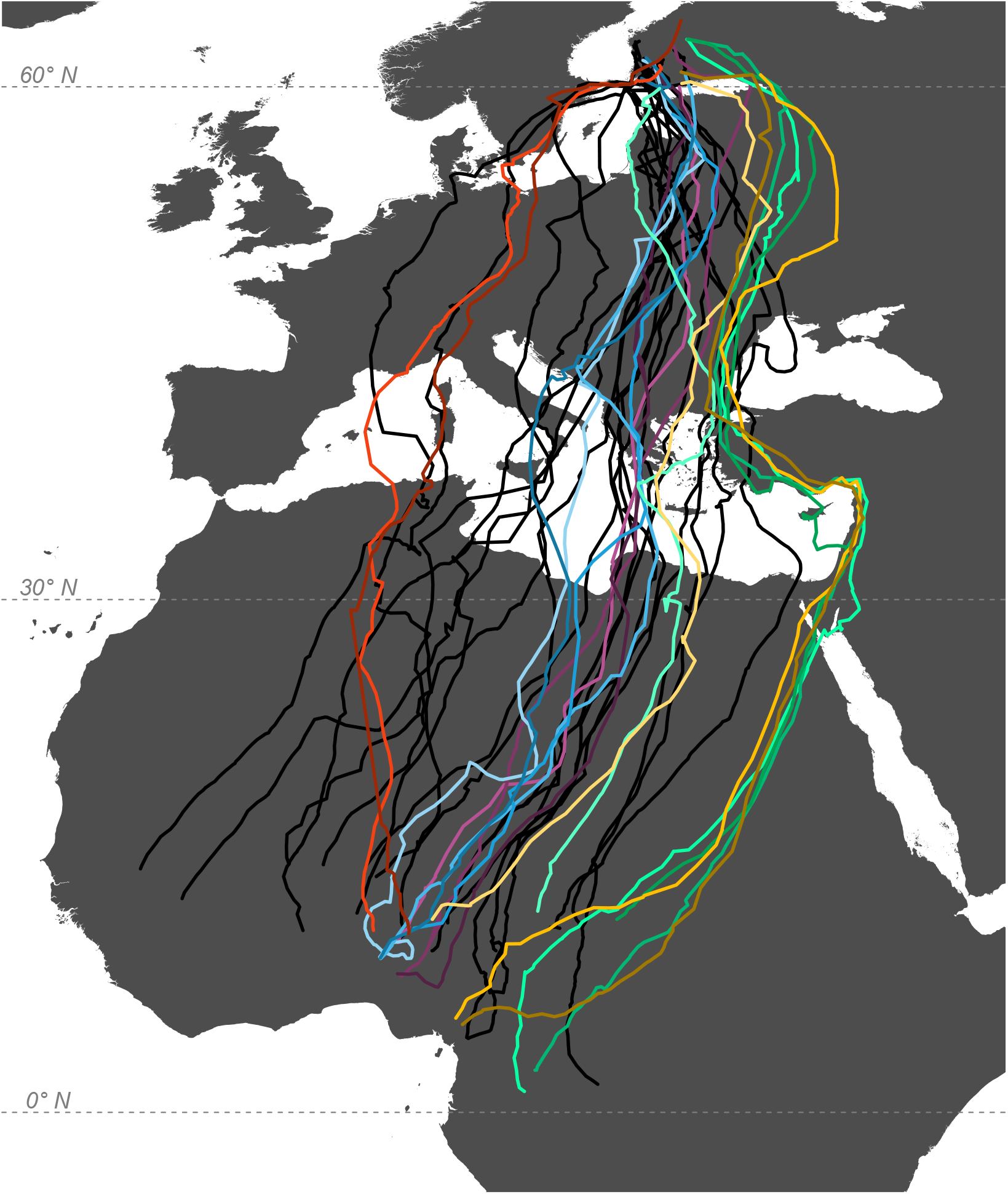
The routes taken by individuals of known age. Routes of the individuals that transmitted only one migration are shown in black, those of the remaining five individuals are shown in colors. The first migrations of multi-year individuals are shown in pale shades that get darker in each consecutive migration.

**Supplemental material 4:**
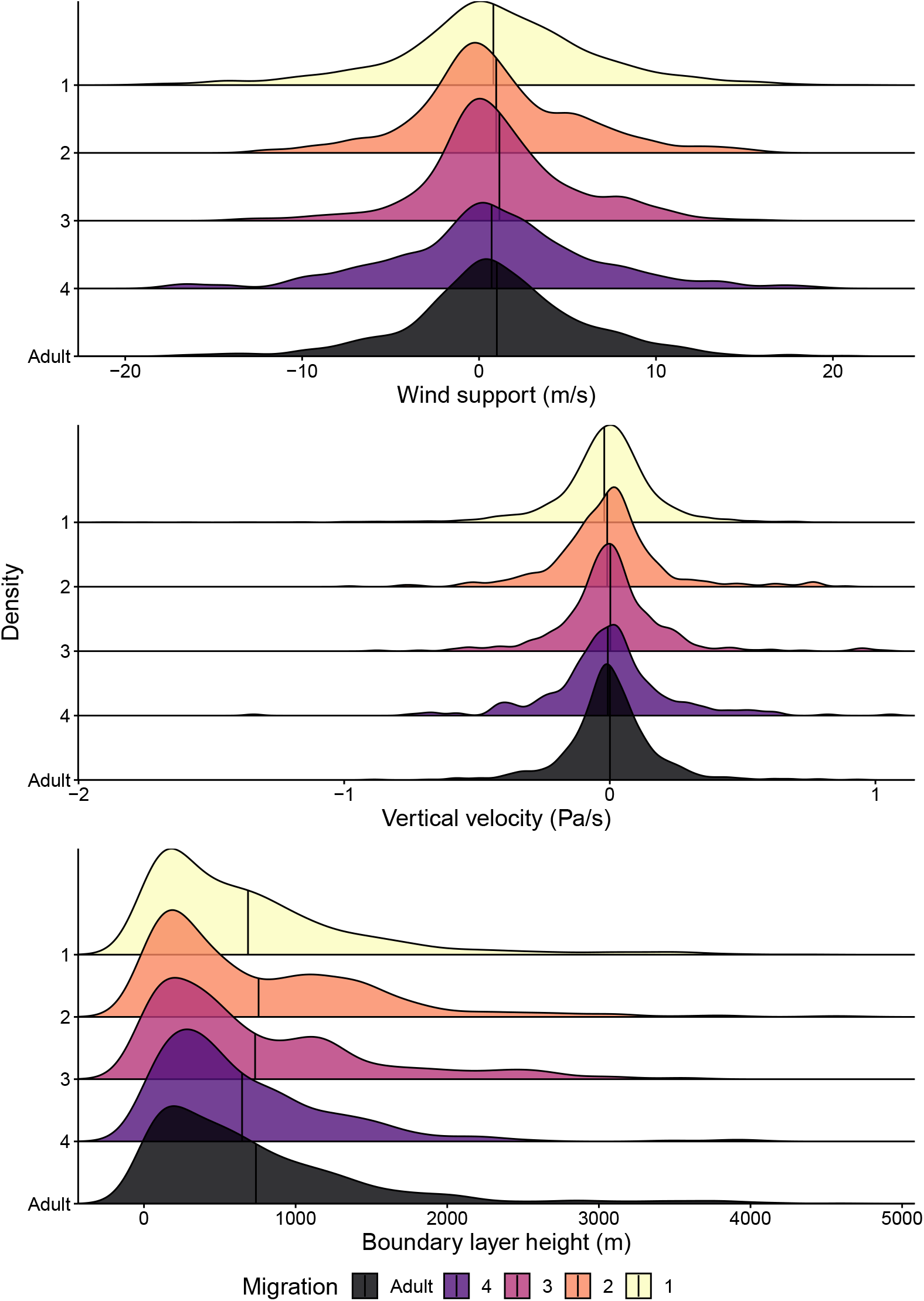
Used atmospheric conditions do not differ among migrations. The distributions of wind support (top), vertical velocity of pressure (middle), and boundary layer height (bottom) for each migration, along with the mean value (black line).

**Supplemental material 5:**
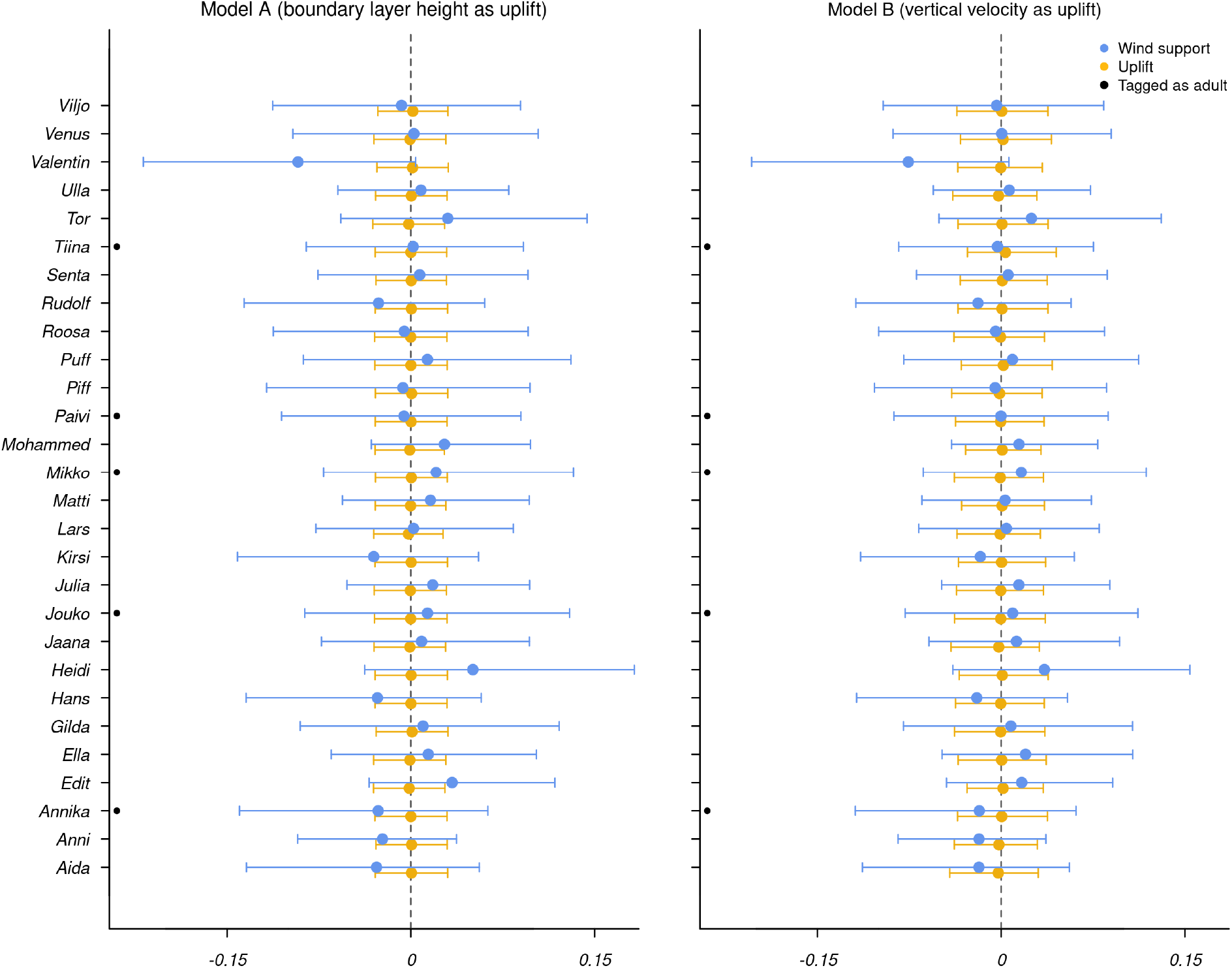
The individual variation in the importance of wind support and uplift for each model.

## Notes

### Competing Interest Statement

The authors have declared no competing interest.

### Summary of Updates

The modeling procedure is changed to include age as a predictor of route selection. The methods, results, figure, and supplemental material are updated to match.

