## Supplemental material 3 (R code) for "Experience does not change the importance of wind support for migratory route selection by a soaring bird"

### Performing analyses using Step Selection Functions

Step-selection functions model animal movement as a series of discrete steps between consecutive locations (Fortin et al. 2005; Fieberg et al. 2021). Each observed step is matched with alternative steps, which allows comparisons of the observed locations to alternative, available ones. Here, we generate these alternative steps and available locations and compare the available airflow.

Step-selections functions define availability with three criteria: the previous location, the time between consecutive locations, and the characteristics of an individual's movement. First, we have to clean the data and process them into tracks. Then, we have to re-sample them so that there are consistent time intervals between locations. Finally, we have to extract the step lengths and turn angles for each individual and generate distributions of what is possible.

Once we have generated the available locations, we can label them with environmental information using Movebank's Env-DATA service (Dodge et al. 2013). The last step is to ask how these environmental factors predict space use, which we do by fitting conditional logistic regressions.

#### 1. Prepare the environment

The first step is to make sure that we have all of the packages and functions that we need. Here, we will use the "amt" package (Signer, Fieberg, and Avgar 2019) to make and process our tracks and the "INLA" package (Lindgren and Rue 2015) to model our data. We need several additional packages for formatting, mapping, and ease.

---

```
#set a working directory and load required packages
setwd("C:/Users/Tess Bronnvik/Desktop/Br-nnvik_honey_buzzard_ssf")
mypath <- paste0(getwd(),"/")
ssf_packs <- c("lubridate", "amt", "move", "mapview", "ggpubr", "survival", "INLA", "ggregplot",
               "magrittr", "sjPlot", "MASS", "modelr", "mosaic", "caret", "viridis", "tidyverse")
lapply(ssf_packs, require, character.only = TRUE)

#import required functions
NCEP.loxodrome.na <- function (lat1, lat2, lon1, lon2) {
  deg2rad <- pi/180
  acot <- function(x) {
    return(atan(1/x))
  }
  lat1 <- deg2rad * lat1
  lat2 <- deg2rad * lat2
```

```r
  lon1 <- deg2rad * lon1
  lon2 <- deg2rad * lon2
  deltaLon <- lon2 - lon1
  pi4 <- pi/4
  Sig1 <- log(tan(pi4 + lat1/2))
  Sig2 <- log(tan(pi4 + lat2/2))
  deltaSig <- Sig2 - Sig1
  if (deltaLon == 0 && deltaSig > 0) {
    head <- 0
  }
  else if (deltaLon == 0 && deltaSig < 0) {
    head <- 180
  }
  else if (deltaSig == 0 && deltaLon > 0) {
    head <- 90
  }
  else if (deltaSig == 0 && deltaLon < 0) {
    head <- 270
  }
  else if (deltaSig < 0 && deltaLon < 0) {
    head <- acot(deltaSig/deltaLon) * 180/pi + 180
  }
  else if (deltaSig < 0 && deltaLon > 0) {
    head <- acot(deltaSig/deltaLon) * 180/pi + 180
  }
  else if (deltaSig > 0 && deltaLon > 0) {
    head <- acot(deltaSig/deltaLon) * 180/pi
  }
  else if (deltaSig > 0 && deltaLon < 0) {
    head <- acot(deltaSig/deltaLon) * 180/pi + 360
  }
  else {
    head <-NA}
  return(head)
}
# functions for calculating wind support from N/S and E/W wind and birds' directions of
# travel
source(paste0(mypath, "wind_support_Kami.R"))
```

---

Next, we set some criteria for building the tracks. The step number determines how many alternative steps are generated for each observed one. The tolerance determines how many minutes around a fix are allowed in order to consider it a next step (eg. a step is the distance traveled in 2 hours +/- 15 minutes). Finally, the CRS is the coordinate reference system, defining the map projection for the mk_track function. Here we have to inform R that our data are measured in degrees rather than in meters.

---

```r
# set criteria for tracks
stepNumber <- 100 # random steps. The number of steps used is arbitrary. The goal is to use
# enough to capture the actual distribution of available environmental conditions without
# running out of computational ability.
```

```r
toleranceLength <- 15 # tolerance in minutes. We use 15 minutes because some of our steps
# are only 1 hour apart and we need to keep the timing precise.
wgs <- CRS("+proj=longlat +datum=WGS84 +no_defs") # map projection
```

---

**2. Prepare data for Movebank's environmental data annotation**

In order to get the data in the right format, we select only the reads from autumn migrations and clean them. We need to remove any duplicated reads and any times when the birds were not moving. We also need to map the data to see whether there are any incomplete records, and to remove those records if we find any. Then we need to separate the data so that individuals that were sampled at different rates are analyzed separately.

---

```r
# retrieve the full data set
full_data <- read.csv(paste0(mypath,"original_data/full_buzz_set.csv"),
                      stringsAsFactors = F, header = T)

# reformat the time stamps
full_data$timestamp <- as.POSIXct(strptime(full_data$dt, format = "%Y-%m-%d %H:%M:%S"),
                                  tz = "UTC")

# select the autumn migrations
all_autumns <- full_data[grep("autumn", full_data$phase),]

# find and remove duplicate observations
doubles <- all_autumns %>% dplyr::select(long, lat, name, timestamp) %>%
  duplicated
sum(doubles) # 59 duplicates
```

```
## [1] 59
```

```r
all_autumns <- all_autumns[doubles != TRUE,]

# get the 204 reads when the birds did not move and remove them from the data
resting_reads <- read.csv(paste0(mypath,"resting.csv"), stringsAsFactors = F)
all_autumns <- all_autumns %>% mutate(id_ts = paste(name,timestamp, sep="_")) %>%
  filter(!id_ts %in% resting_reads$id_ts)

# order all the data by timestamp
all_autumns <- all_autumns %>%
  arrange(timestamp)

# create a unique identifier for each migratory journey
all_autumns$id_year <- paste(all_autumns$name, all_autumns$yr, sep="_")

# make simple plots of lat/long to check for outliers
ggplot(all_autumns, aes(x=long, y=lat, color=as.factor(name))) + geom_point() +
  theme(legend.position = "none")
```

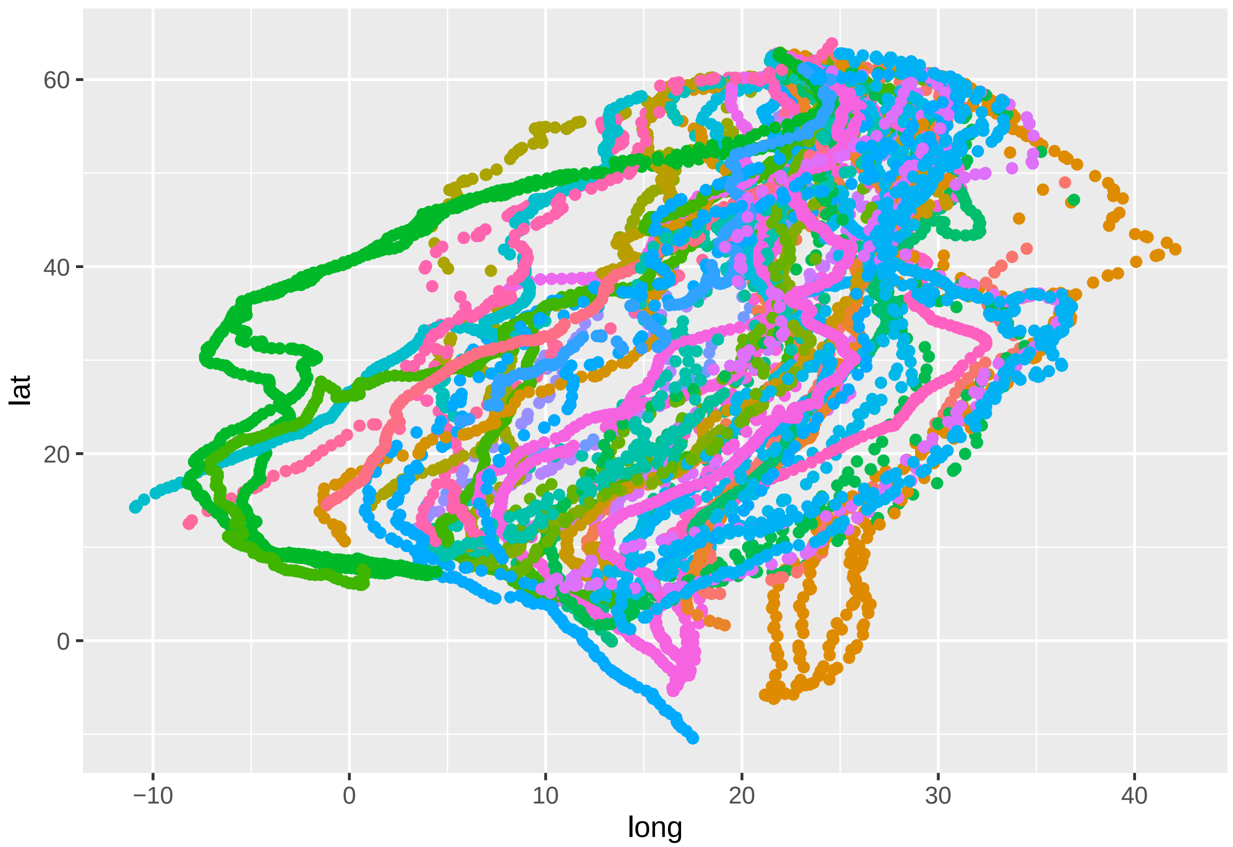


```
ggplot(all_autumns, aes(x=long, y=lat)) + geom_point()+ facet_wrap(~name, scales="free")
```

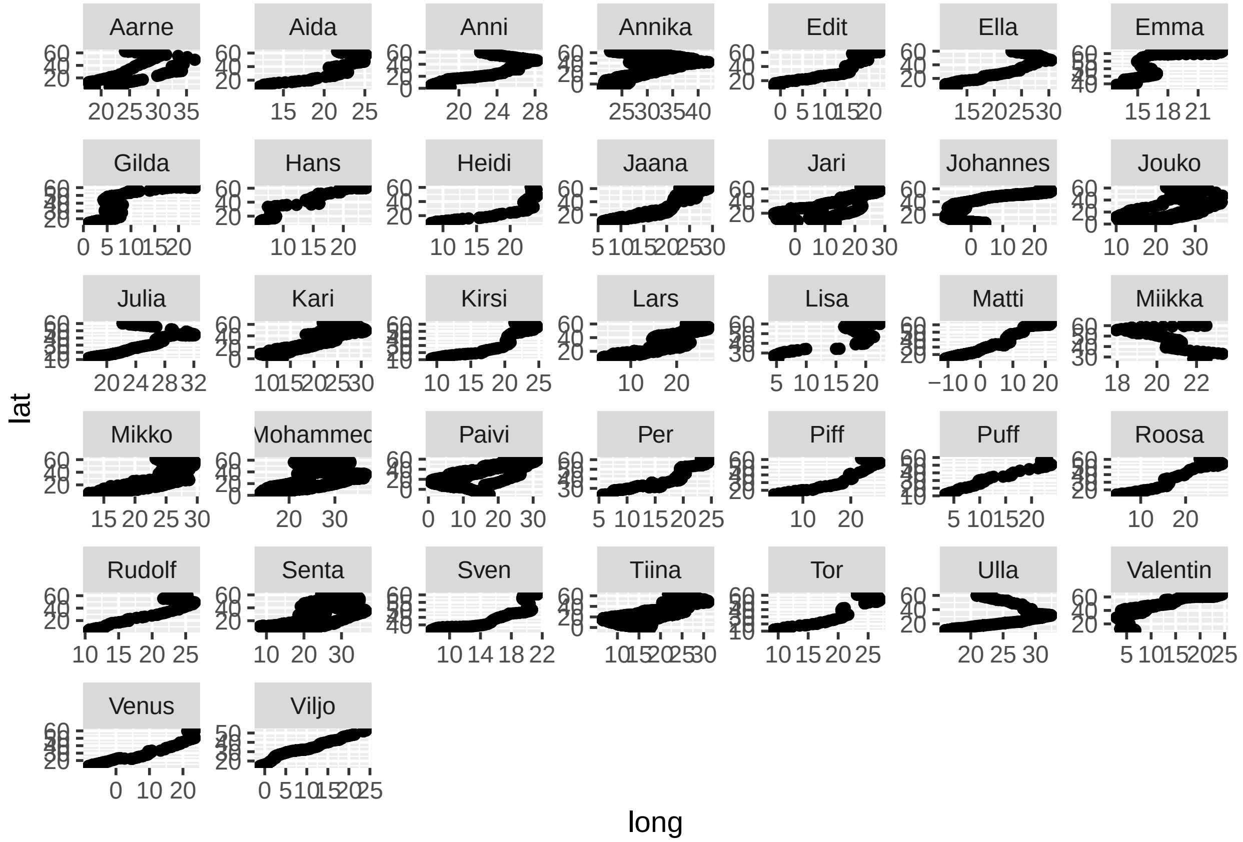


```
# make a map of individual tracks
data_sp <- all_autumns # store the data, then create a spatial object for plotting
coordinates(data_sp) <- ~ long + lat # set coordinates
proj4string(data_sp) <- wgs # set projection
mapView(data_sp, zcol = "id_year", burst = F, cex = 3, color = rainbow) # plot on a map
```

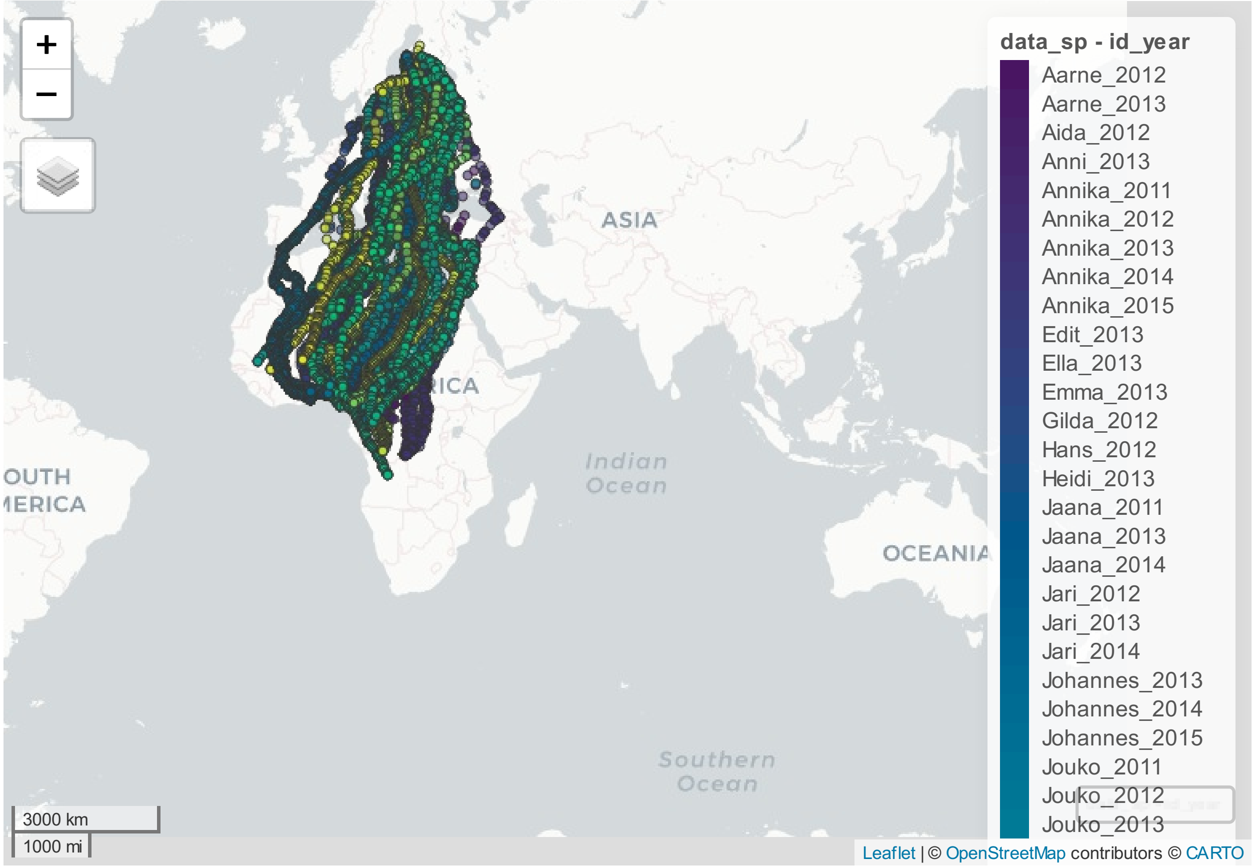


```
# we don't know the amount of experience that birds tagged as adults have
# collect the journeys for which experience can't be safely guessed
unk_age <- c("Annika_2011", "Annika_2012", "Annika_2013", "Jouko_2011",  "Jouko_2012",
             "Jouko_2013", "Mikko_2011", "Mikko_2012", "Mikko_2013",  "Paivi_2013",
             "Paivi_2014", "Paivi_2015", "Tiina_2011",  "Tiina_2012", "Tiina_2013")

# remove birds of unknown age with fewer than 4 tracked autumn migrations
all_autumns <- all_autumns %>% filter(name != "Aarne" &
                        name != "Jari" &
                        name != "Johannes" &
                        name != "Kari" &
                # disregard the data from early migrations by birds of unknown age
                       !id_year %in% unk_age &
                # some individuals do not transmit their first trip
                        name != "Emma" &
                        name != "Lisa" &
                        name != "Miikka" &
                        name != "Per" &
                        name != "Sven" &
                # others are lost en route later in life and must also be removed
                        id_year != "Annika_2015" &
                        id_year != "Jouko_2015" &
                        id_year != "Paivi_2017" &
                        id_year != "Senta_2017")

# because the loggers have different average sampling rates,
```

```
# the data need to be re-sampled at different rates. Label them with these rates
# in minutes so that each can be processed individually.
all_autumns$sample_rate <- NA

# the individuals with 1 hour sampling rates
all_autumns$sample_rate[which(all_autumns$name == "Anni")] <- 60

# the individuals with 2 hour sampling rates
all_autumns$sample_rate[which(all_autumns$name == "Edit" | all_autumns$name == "Julia" |
                              all_autumns$name == "Matti"| all_autumns$name == "Ulla" |
                              all_autumns$name == "Aida" | all_autumns$name == "Ella" |
                              all_autumns$name == "Heidi" | all_autumns$name == "Kirsi" |
                              all_autumns$name == "Gilda" |
                              all_autumns$name == "Valentin")] <- 120

# the individuals with 3 hour sampling rates
all_autumns$sample_rate[which(all_autumns$name == "Mohammed")] <- 180

# the individuals with 4 hour sampling rates
all_autumns$sample_rate[which(all_autumns$name == "Annika" | all_autumns$name == "Jaana" |
                              all_autumns$name == "Lars"| all_autumns$name == "Piff"|
                              all_autumns$name == "Puff" | all_autumns$name == "Roosa"|
                              all_autumns$name == "Senta"| all_autumns$name == "Tor" |
                              all_autumns$name == "Tiina" |all_autumns$name == "Jouko" |
                              all_autumns$name == "Mikko" |all_autumns$name == "Paivi" |
                              all_autumns$name == "Hans" | all_autumns$name == "Venus" |
                              all_autumns$name == "Rudolf")] <- 240
```

---

Using the amt package, we can create track objects. These are then re-sampled to a given step length. Steps have to be of consistent times because the parameters of the models are scale dependent and will vary with $\Delta$t. By re-sampling, we burst the tracks. A burst is a track segment containing fixes separated by the given step length. We can then create randomized locations as part of these bursts.

---

```
# build tracks at each rate

autumn_track <- data.frame()

for (i in sort(unique(all_autumns$sample_rate))) {
  # select data of the given sampling rate
  temp_rdata <- all_autumns[which(all_autumns$sample_rate == i),]
  for (j in sort(unique(temp_rdata$id_year))) {
  print(j)
  # select data for the given track
  temp_idata <- temp_rdata[which(temp_rdata$id_year == j),]
  # make the track
  trk <- mk_track(temp_idata,.x=long, .y=lat, .t=timestamp, id = name, crs = wgs)
  # resample to a consistent time between steps
  trk <- track_resample(trk, rate = minutes(i), tolerance = minutes(toleranceLength))
  # remove bursts with fewer than three re-locations so that turn angle can be calculated
```

```
  trk <- filter_min_n_burst(trk, 3)
  # convert track to a step representation, and calculate sl_, direction, and ta_
  burst <- steps_by_burst(trk, keep_cols = "start")#, lonlat = T)
  # create random steps using fitted gamma and von Mises distributions (as suggested by
  # previous studies) and append
  rnd_stps <- burst %>% random_steps(n_control = stepNumber)
  # save
  autumn_track <- rbind(autumn_track, rnd_stps)
  # and signal
  print(paste0("Successfully created random steps for track ", j, "."))
  }
}

# compare the step lengths and turn angles for observed and random steps
obs_sl <- autumn_track[autumn_track$case_ == TRUE,] %>%
  ggplot(aes(sl_, fill = factor(id))) + ggtitle("Observed") +
  geom_density(alpha = 0.4) +
  labs(x = "Step length (degrees)", y = "Density") +
  theme_minimal()
obs_ta <- autumn_track[autumn_track$case_ == TRUE,] %>%
  ggplot(aes(ta_, fill = factor(id))) + ggtitle("Observed") +
  geom_density(alpha = 0.4) +
  labs(x = "Turn angle (radians)", y = "Density") +
  theme_minimal()
rand_sl <- autumn_track[autumn_track$case_ == FALSE,] %>%
  ggplot(aes(sl_, fill = factor(id))) + ggtitle("Alternative") +
  geom_density(alpha = 0.4) +
  labs(x = "Step length (degrees)", y = "Density") +
  theme_minimal()
rand_ta <- autumn_track[autumn_track$case_ == FALSE,] %>%
  ggplot(aes(ta_, fill = factor(id))) + ggtitle("Alternative") +
  geom_density(alpha = 0.4) +
  labs(x = "Turn angle (radians)", y = "Density") +
  theme_minimal()
ggarrange(obs_sl, rand_sl, obs_ta, rand_ta, ncol=2, nrow=2, legend = "none")
```

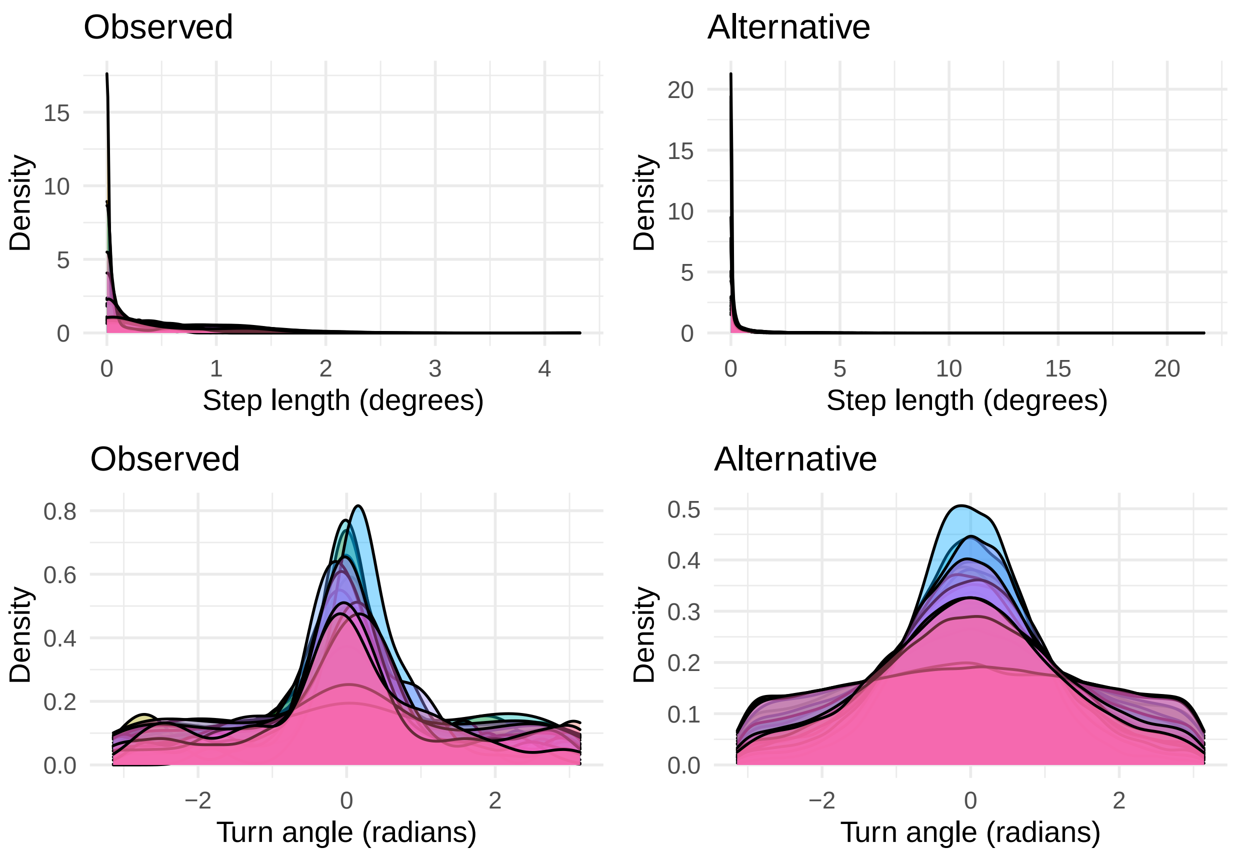


---

In order to calculate the wind support component of the north/south and east/west winds that we will get from Movebank annotation, we need to calculate the direction of movement (loxodrome angle of travel) between locations. This is best done before annotation to avoid truncation of smaller values in the locations during file export.

---

```
# calculate directions of movement between locations for each step
autumn_track <- autumn_track %>%
  rowwise() %>%
  mutate(heading = NCEP.loxodrome.na(y1_, y2_, x1_, x2_)) %>%
  ungroup()
```

---

Now we only need to conform to column name and file size specifications, then export the prepared data and send it to the Env-DATA service.

---

```
# select the end point for each step
names(autumn_track)[c(2,4)] <-c("location-long", "location-lat")
```

```
# select the start time for each step
autumn_track$timestamp <- autumn_track$t1_
autumn_track$timestamp <- paste0(autumn_track$timestamp,".000" )

# export the data for Movebank annotation
write.csv(autumn_track, paste0("HB_", Sys.Date(), ".csv"), row.names = FALSE)
```

---

#### 3. Prepare data for modeling

For these step-selection analyses, we use integrated nested Laplace approximation (INLA) to predict whether a location is used or unused based on environmental conditions and age. Once we have our files returned from Movebank's Env-DATA service, we can process the annotated data for modeling. Using the directions we calculated above, we can derive wind support (m/s) from the wind data. Our second variable of interest is uplift. We use two proxies for uplift: 1) planetary boundary layer height (m), which is higher where rising air pushes it upward (ECMWF 2018a), and 2) vertical velocity of pressure (Pa/s), which is a measure of rising air for which negative values indicate upward motion (ECMWF 2018b). Our third variable is the interaction of wind support, uplift, and age.

---

```
annotated_data <- read.csv(paste0(mypath,
                           "annotations/HB_2022-04-29-526598093103552488.csv"),
                           stringsAsFactors = F, header = T)  %>%
  # rename the columns to be more useful
  rename(lon1 = x1_,
         lat1 = y1_,
         lon2 = location.long,
         lat2 = location.lat,
         u_wind = ECMWF.ERA5.PL.U.Wind,
         v_wind = ECMWF.ERA5.PL.V.Wind,
         vertical_pressure = ECMWF.ERA5.PL.Pressure.Vertical.Velocity,
         BLH = ECMWF.ERA5.SL.Boundary.Layer.Height) %>%
  # then reformat the time and logical, and calculate wind support
  mutate(timestamp = as.POSIXct(strptime(timestamp, format = "%Y-%m-%d %H:%M:%S"),
                                tz = "UTC"),
         year = format(as.POSIXct(timestamp,format="%Y-%m-%d %H:%M:%S"),"%Y"),
         id_year = paste(id,year,sep="_"),
         case_ = as.numeric(case_),
         tail = wind_support(u_wind, v_wind, heading))

# label the data with life stage and migration information
annotated_data$stage <- NA
annotated_data$stage[which(annotated_data$id == "Annika" |
                             annotated_data$id == "Jouko" |
                             annotated_data$id == "Mikko" |
                             annotated_data$id == "Paivi" |
                             annotated_data$id == "Tiina" )] <- "adult"
annotated_data$stage[which(is.na(annotated_data$stage))] <- "juvenile"

annotated_data$migration <- NA # create an empty column to store values
```

```
annotated_data$migration[which(annotated_data$id_year == "Jaana_2013" | # add year 2 info
                                 annotated_data$id_year == "Lars_2013" |
                                 annotated_data$id_year == "Mohammed_2018" |
                                 annotated_data$id_year == "Senta_2015" |
                                 annotated_data$id_year == "Valentin_2015")] <- "2"

annotated_data$migration[which(annotated_data$id_year == "Jaana_2014" | # add year 3 info
                                 annotated_data$id_year == "Lars_2014" |
                                 annotated_data$id_year == "Mohammed_2019" |
                                 annotated_data$id_year == "Senta_2016")] <- "3"

annotated_data$migration[which(annotated_data$id_year == "Lars_2015" | # add year 4 info
                                 annotated_data$id_year == "Mohammed_2020")] <- "4"

annotated_data$migration[which(annotated_data$id_year == "Annika_2014" | # add year 5 info
                            annotated_data$id_year == "Jouko_2014" |
                            annotated_data$id_year == "Mikko_2014" |
                            annotated_data$id_year == "Mikko_2015" |
                            annotated_data$id_year == "Paivi_2016" |
                            annotated_data$id_year == "Tiina_2014" |
                            annotated_data$id_year == "Tiina_2015")] <- "Adult"
annotated_data$migration[which(is.na(annotated_data$migration))] <- "1" # everything else is year 1
```

---

```
# compare the distributions of used environmental conditions among ages
annotated_data$migration <- factor(annotated_data$migration, levels = c("Adult", 4, 3, 2, 1))

ws <- annotated_data[annotated_data$case_ == TRUE,] %>%
    ggplot(aes(tail, migration, fill = migration, color = NULL)) +
    geom_density_ridges2(alpha = 0.8, quantile_lines=TRUE,
                      quantile_fun=function(x,...)mean(x)) +
    labs(x = "Wind support (m/s)", y = "", fill = "Migration") +
    scale_fill_viridis(discrete = TRUE, option = "A") +
    coord_cartesian(expand = F) +
    theme_minimal() +
    theme(legend.position = "none", text = element_text(size=15),
        axis.text.x = element_text(colour = "black"),
        axis.text.y = element_text(colour = "black"),
        panel.grid.major = element_blank(),
        panel.grid.minor = element_blank(),
        strip.text.x = element_text(size = 20),
        axis.line = element_line(colour = "black"),
        axis.ticks = element_line(colour = "black"))
vv <- annotated_data[annotated_data$case_ == TRUE,] %>%
    ggplot(aes(vertical_pressure, migration, fill = migration)) +
    geom_density_ridges2(alpha = 0.8,quantile_lines=TRUE,
                      quantile_fun=function(x,...)mean(x)) +
    labs(x = "Vertical velocity (Pa/s)", y = "Density", fill = "Migration") +
    scale_fill_viridis(discrete = TRUE, option = "A") +
    coord_cartesian(expand = F)+
    theme_minimal() +
    theme(legend.position = "none", text = element_text(size=15),
        axis.text.x = element_text(colour = "black"),
```

```
        axis.text.y = element_text(colour = "black"),
        panel.grid.major = element_blank(),
        panel.grid.minor = element_blank(),
        strip.text.x = element_text(size = 20),
        axis.line = element_line(colour = "black"),
        axis.ticks = element_line(colour = "black"))
bl <- annotated_data[annotated_data$case_ == TRUE,] %>%
    ggplot(aes(BLH, migration, fill = migration)) +
    geom_density_ridges2(alpha = 0.8, quantile_lines=TRUE,
                  quantile_fun=function(x,...)mean(x)) +
    labs(x = "Boundary layer height (m)", y = "", fill = "Migration") +
    scale_fill_viridis(discrete = TRUE, option = "A") +
    coord_cartesian(expand = F)+
    theme_minimal() +
    theme(legend.position = "none", text = element_text(size=15),
        axis.text.x = element_text(colour = "black"),
        axis.text.y = element_text(colour = "black"),
        panel.grid.major = element_blank(),
        panel.grid.minor = element_blank(),
        strip.text.x = element_text(size = 20),
        axis.line = element_line(colour = "black"),
        axis.ticks = element_line(colour = "black"))
ggarrange(ws, vv, bl, ncol = 1, nrow = 3, common.legend = T, legend = "bottom")
```

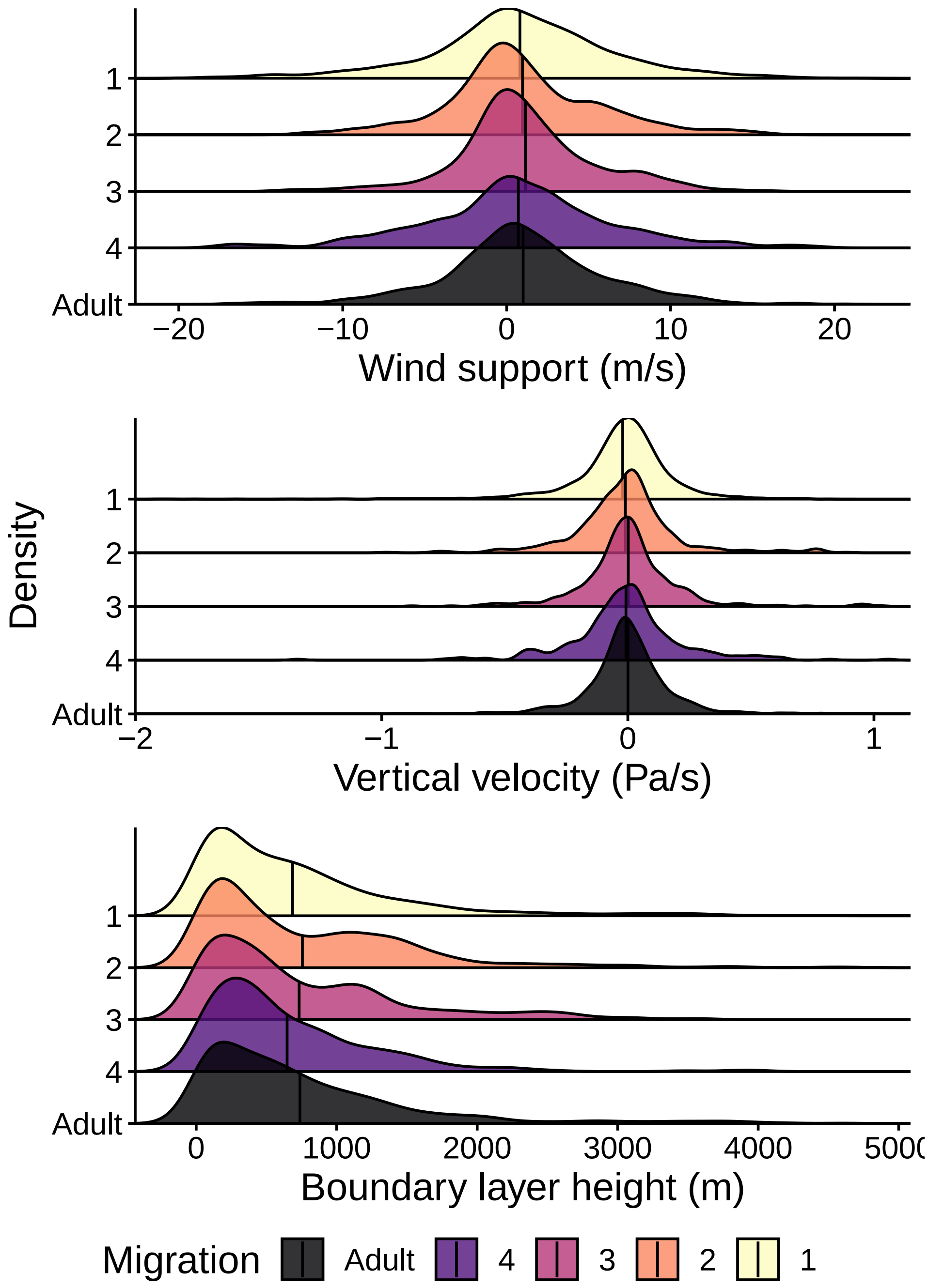


---

##### 4. Fit the models

To evaluate the effects of wind support, uplift, and age on route selection, we fit integrated nested Laplace approximations (Muff, Signer, and Fieberg 2020). This is a recent alternative to Markov chain Monte Carlo that focuses on estimating individual posterior marginals of the model parameters.

---

```
#add columns for ID and year
all_data <- annotated_data %>%
  mutate(migration = as.numeric(ifelse(migration == "Adult", 5, migration)), #create a numeric variable
         id1 = factor(id),
         id2 = factor(id),
         yr1 = factor(year),
         yr2 = factor(year))

all_data <- all_data %>%
  drop_na(tail) %>%
  mutate_at(c("tail", "BLH", "migration" , "vertical_pressure"),
            list(z = ~(scale(.)))) %>%
  rename(stratum = step_id_)

## Build the models:

#model formula
formula_i <- case_ ~ -1 + tail_z * BLH_z * migration_z +
  f(stratum, model = "iid",
    hyper = list(theta = list(initial = log(1e-6),fixed = T))) +
  f(id1, BLH_z, model = "iid",
    hyper=list(theta=list(initial=log(1),fixed=F,prior="pc.prec",param=c(3,0.05)))) +
  f(id2, tail_z,  model = "iid",
    hyper=list(theta=list(initial=log(1),fixed=F,prior="pc.prec",param=c(3,0.05))))

formula_v <- case_ ~ -1 + tail_z * vertical_pressure_z * migration_z +
  f(stratum, model = "iid",
    hyper = list(theta = list(initial = log(1e-6),fixed = T))) +
  f(id1, vertical_pressure_z, model = "iid",
    hyper=list(theta=list(initial=log(1),fixed=F,prior="pc.prec",param=c(3,0.05)))) +
  f(id2, tail_z,  model = "iid",
    hyper=list(theta=list(initial=log(1),fixed=F,prior="pc.prec",param=c(3,0.05))))

all_data <- readRDS("inla_input.rds")

mean.beta <- 0
prec.beta <- 1e-4


# record the start time to check how long the model requires to run
(b <- Sys.time())
#model without missing values
M_v <- inla(formula_v, family = "Poisson",
                 control.fixed = list(
                   mean = mean.beta,
                   prec = list(default = prec.beta)),
                 data = all_data,
                 num.threads = 10,
            # this means that NA values will be predicted.
                 control.predictor = list(compute = TRUE),
                 control.compute = list(openmp.strategy = "huge", config = TRUE, cpo = T))
# check the run time
Sys.time() - b
```

```
#extract CPO and Marginal likelihood values
mean(M_v$cpo$cpo) #model with vertical velocity and random effect for individual id
# 0.9799288

M_v$mlik
# integration -38533.82
#Gaussian -38534.20

mean(M_i$cpo$cpo) #model with BLH and random effect for individual id
#0.9799288

M_i$mlik
# integration : -38518.27
#Gaussian : -38517.73
```

---

**5. Extract information from model output for plotting**

Original code by Virgilio Gomez-Rubio (Gómez-Rubio 2020).

---

```
#extract info for individual variation plots. Done separately for each model
tab_blh <- data.frame(ID = as.factor(M_i$summary.random$id1$ID),
                      mean = M_i$summary.random$id1$mean,
                      IClower = M_i$summary.random$id1[, 4],
                      ICupper = M_i$summary.random$id1[, 6])

head(tab_blh_M_i)
```

```
##       ID          mean     IClower    ICupper
## 1   Aida  5.209592e-04 -0.02908335 0.03014164
## 2   Anni  5.989837e-04 -0.02845070 0.02964099
## 3 Annika  8.124335e-05 -0.02938873 0.02953937
## 4   Edit -1.258232e-03 -0.03040362 0.02781964
## 5   Ella -7.936469e-04 -0.03031122 0.02870819
## 6  Gilda  1.017997e-03 -0.02832759 0.03041957
```

```
tab_wspt <- data.frame(ID = as.factor(M_i$summary.random$id2$ID),
                       mean = M_i$summary.random$id2$mean,
                       IClower = M_i$summary.random$id2[, 4],
                       ICupper = M_i$summary.random$id2[, 6])

#saveRDS(tab_vv, file = "tab_vv_M_v.rds")
#saveRDS(tab_wspt, file = "tab_wspt_M_v.rds")


#saveRDS(tab_blh, file = "tab_blh_M_i.rds")
#saveRDS(tab_wspt, file = "tab_wspt_M_i.rds")
```

```r
# extract info for coefficient plots

# posterior means of coefficients
graph <- as.data.frame(summary(M_i)$fixed)
colnames(graph)[which(colnames(graph)%in%c("0.025quant","0.975quant"))]<-c("Lower","Upper")
colnames(graph)[which(colnames(graph)%in%c("0.05quant","0.95quant"))]<-c("Lower","Upper")
colnames(graph)[which(colnames(graph)%in%c("mean"))]<-c("Estimate")

#graph$Model<-i
graph$Factor <- rownames(graph)


#saveRDS(graph, file = "graph_M_v.rds")
#saveRDS(graph, file = "graph_M_i.rds")

# the results for Model A (boundary layer height)
head(graph_M_i)

##                     Estimate    sd  Lower 0.5quant Upper mode kld
## tail_z                 0.095 0.018  0.059    0.095 0.131   NA   0
## BLH_z                 -0.020 0.014 -0.049   -0.020 0.008   NA   0
## migration_z            0.001 0.014 -0.026    0.001 0.027   NA   0
## tail_z:BLH_z           0.090 0.017  0.058    0.090 0.123   NA   0
## tail_z:migration_z     0.012 0.016 -0.020    0.013 0.044   NA   0
## BLH_z:migration_z      0.009 0.014 -0.018    0.009 0.036   NA   0
##                                 Factor
## tail_z                          tail_z
## BLH_z                            BLH_z
## migration_z                migration_z
## tail_z:BLH_z              tail_z:BLH_z
## tail_z:migration_z tail_z:migration_z
## BLH_z:migration_z   BLH_z:migration_z

# the results for Model B (vertical velocity of pressure)
head(graph_M_v)

##                                  Estimate    sd  Lower 0.5quant Upper mode kld
## tail_z                              0.108 0.017  0.074    0.108 0.142   NA   0
## vertical_pressure_z                -0.013 0.014 -0.041   -0.013 0.015   NA   0
## migration_z                         0.000 0.014 -0.026    0.000 0.027   NA   0
## tail_z:vertical_pressure_z          0.008 0.010 -0.011    0.008 0.028   NA   0
## tail_z:migration_z                  0.011 0.016 -0.020    0.011 0.042   NA   0
## vertical_pressure_z:migration_z     0.000 0.014 -0.029    0.000 0.028   NA   0
##                                                           Factor
## tail_z                                                    tail_z
## vertical_pressure_z                          vertical_pressure_z
## migration_z                                          migration_z
## tail_z:vertical_pressure_z            tail_z:vertical_pressure_z
## tail_z:migration_z                            tail_z:migration_z
## vertical_pressure_z:migration_z vertical_pressure_z:migration_z
```

#### 6. Plot the results

---

```
gr_blh <- readRDS(paste0(mypath, "graph_M_i.rds"))
gr_vv <- readRDS(paste0(mypath, "graph_M_v.rds"))


#X11(width = 8.4, height = 3)
png(paste0(mypath, "coeff_plot.png"), units = "in", width = 8.4, height = 3, res = 300)

par(mfrow = c(1,2),
    cex = 0.7,
    oma = c(0,14,0,0),
    mar = c(3, 1, 3, 1),
    bty = "l",
    #font = 3,
    font.axis = 3
)


for(graph in list(gr_blh,gr_vv)){


VarOrder <- rev(unique(graph$Factor))
VarNames <- VarOrder

graph$Factor <- factor(graph$Factor, levels = VarOrder)
levels(graph$Factor) <- VarNames

min <- min(graph$Lower,na.rm = T)
max <- max(graph$Upper,na.rm = T)

graph$Factor_n <- as.numeric(graph$Factor)


plot(0, type = "n", labels = FALSE, tck = 0, xlim = c(-0.05,0.14), ylim = c(0.7,7.3), xlab = "Estimate"

#add vertical line for zero
abline(v = 0, col = "grey30",lty = 2)
#add points and error bars
points(graph$Estimate, graph$Factor_n, col = "cornflowerblue", pch = 20, cex = 2)
arrows(graph$Lower, graph$Factor_n,
       graph$Upper, graph$Factor_n,
       #angle of 90 to make the arrow head as straight as a line
       col = "cornflowerblue", code = 3, length = 0.03, angle = 90, lwd = 2)

#add axes
axis(side= 1, at = c(-0.04, 0, 0.04, 0.08,0.12), labels =  c(-0.04, 0, 0.04, 0.08, 0.12),
     tick=T ,col = NA, col.ticks = 1, tck =-.015)

if("BLH_z" %in% graph$Factor){
axis(side = 2, at = c(1:7),
     labels = c("Wind support : Uplift : Migration year ", "Uplift : Migration year", "Wind support : Mi
```

```
                "Wind support"),
     tick=T ,col = NA, col.ticks = 1, # NULL would mean to use the defult color specified by "fg" in pa
     tck=-.015 , #tick marks smaller than default by this proportion
     las=2) # text perpendicular to axis label
  mtext("Model A (boundary layer height as uplift)", side = 3, cex = 0.8, line = 1)

} else {
  axis(side = 2, at = c(1:7),
       labels = c("","", "", "", "", "", ""),
       tick=T ,col = NA, col.ticks = 1, # NULL would mean to use the defult color specified by "fg" in p
       tck=-.015 , #tick marks smaller than default by this proportion
       las=2) # text perpendicular to axis label

  mtext("Model B (vertical velocity as uplift)", side = 3, cex = 0.75, line = 1)
}

}
```

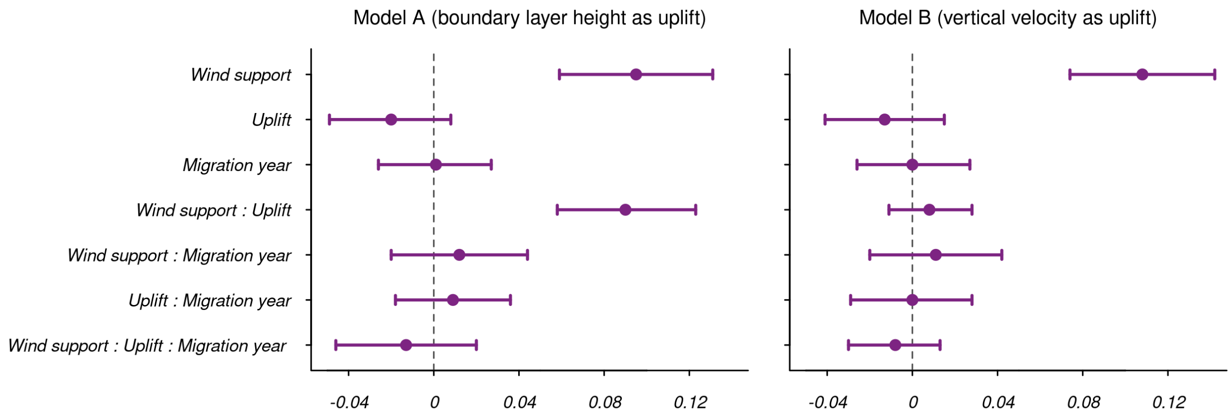


Figure 1: The importance of wind support, uplift, and their interaction to route selection over migrations. Posterior means (centred and scaled) and 95% credible intervals for the fixed effects in the INLA models are shown.

```
dev.off()

age_id <- annotated_data %>%
  group_by(id) %>%
  summarize(age = as.character(head(category,1))) %>% #extract adult vs first year data
  arrange(age) %>%
  as.data.frame()

#re-order names based on age
VarOrder <- rev(unique(graph$Factor))
VarNames <- VarOrder

graph$Factor <- factor(graph$Factor, levels = VarOrder)
levels(graph$Factor) <- VarNames

#plot
```

```
#X11(width = 9, height = 7)

png(paste0(mypath, "ind_var.png"), units = "in", width = 9, height = 7, res = 300)

par(mfrow = c(1,2),
    bty="n",
    cex = 0.7,
    oma = c(0,3.5,0,3),
    mar = c(3, 2, 3,1),
    font.axis = 3
)

for(var in c("i","v")){ #all files with i are related to the model with blh. all with v are related to v

  variables <- str_sub(list.files(paste0(mypath, "Hester_HB/"), pattern = paste0("^tab.*", var), full.na

  files <- list.files(paste0(mypath, "Hester_HB/"), pattern = paste0("^tab.*", var), full.names = T) %>%
    lapply(readRDS)

  names(files) <- variables

  plot(0, bty = "l", labels = FALSE, tck = 0, xlim = c(-0.23,0.17), ylim = c(0,29.5), xlab = "", ylab =
  #add vertical line for zero
  abline(v = 0, col = "grey30",lty = 2)


  points(files[[1]]$mean, as.numeric(files[[1]]$ID) - 0.2, col = "darkgoldenrod2", pch = 19, cex = 1.3)
  arrows(files[[1]]$IClower, as.numeric(files[[1]]$ID) - 0.2,
         files[[1]]$ICupper, as.numeric(files[[1]]$ID) - 0.2,
         col = "darkgoldenrod2", code = 3, length = 0.03, angle = 90, lwd = 1) #angle of 90 to make the

  points(files[[2]]$mean, as.numeric(files[[2]]$ID) , col = "cornflowerblue", pch = 19, cex = 1.3)
  arrows(files[[2]]$IClower, as.numeric(files[[2]]$ID) ,
         files[[2]]$ICupper, as.numeric(files[[2]]$ID) ,
         col = "cornflowerblue", code = 3, length = 0.03, angle = 90, lwd = 1)

  axis(side= 1, at = c(-0.15,0,0.15), labels = c(-0.15,0,0.15),
       tick=T ,col = NA, col.ticks = 1, tck=-.015)

  #add title
  if( var == "i"){
    axis(side= 2, at= c(1:28),
         labels = files[[2]]$ID,
         tick = T ,col = NA, col.ticks = 1,
         tck = -.015 ,
         las = 2)

    mtext("Model A (boundary layer height as uplift)", side = 3, cex = 0.8, line = 0.5)

    } else {
    mtext("Model B (vertical velocity as uplift)", side = 3, cex = 0.75, line = 0.5)
      axis(side= 2, at= c(1:28),
           labels =  NA,
           tick = T ,col = NA, col.ticks = 1,
```

```
        tck = -.015 ,
        las = 2)

    #add legend
    legend("topright", inset=c(-0.01,0), legend = c("Wind support", "Uplift", "Tagged as adult"),
           col = c("cornflowerblue","darkgoldenrod1", "black"),
           pch = 19, bg="white",bty="n", cex = 0.9)
    }

  points(x =rep(-.24,5), y = which(files[[2]]$ID %in% age_id[age_id$age == "Adult","id"]), pch = 20,cex

}
```

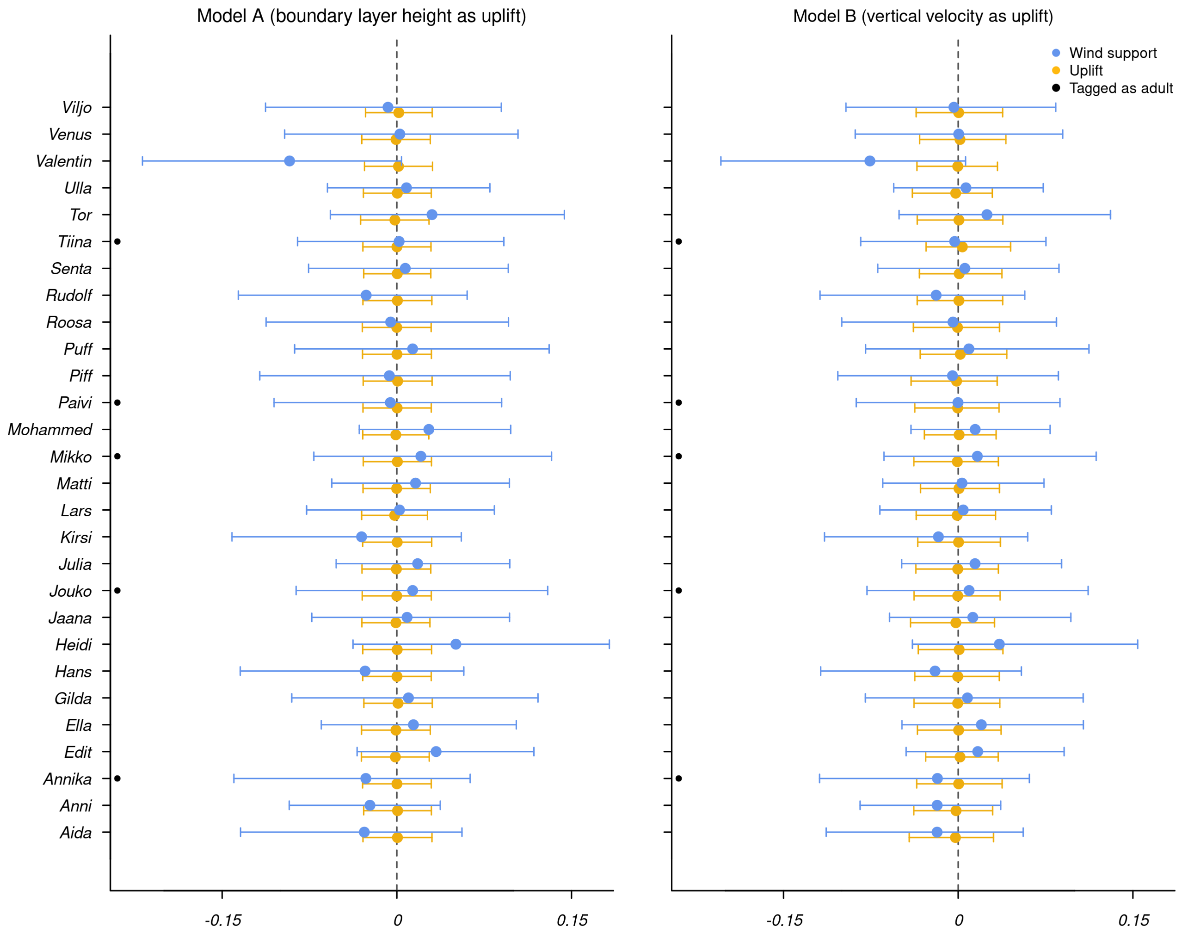


Figure 2: The individual variation in the importance of wind support and uplift for each model.

```
dev.off()
```

ECMWF. 2018b. "GRIB Parameter Database."

———. 2018a. "GRIB Parameter Database."

Fieberg, John, Johannes Signer, Brian Smith, and Tal Avgar. 2021. "A 'How to'guide for Interpreting Parameters in Habitat-Selection Analyses." *Journal of Animal Ecology* 90 (5): 1027–43.

Fortin, Daniel, Hawthorne L Beyer, Mark S Boyce, Douglas W Smith, Thierry Duchesne, and Julie S Mao. 2005. "Wolves influence elk movements: behavior shapes a trophic cascade in Yellowstone National Park." *Ecology* 86 (5): 1320–30. https://doi.org/10.1890/04-0953.

Gómez-Rubio, Virgilio. 2020. *Bayesian Inference with INLA*. CRC Press.

Lindgren, Finn, and Håvard Rue. 2015. "Bayesian Spatial Modelling with r-INLA." *Journal of Statistical Software* 63: 1–25.

Muff, Stefanie, Johannes Signer, and John Fieberg. 2020. "Accounting for Individual-Specific Variation in Habitat-Selection Studies: Efficient Estimation of Mixed-Effects Models Using Bayesian or Frequentist Computation." *Journal of Animal Ecology* 89 (1): 80–92.

Signer, Johannes, John Fieberg, and Tal Avgar. 2019. "Animal movement tools (amt): R package for managing tracking data and conducting habitat selection analyses." *Ecology and Evolution* 9: 880–90. https://doi.org/10.1002/ece3.4823.
